# Prevalence of Erythromycin Resistant *emm92*-type Invasive Group A Streptococcal Infections in West Virginia, United States, 2021-2023

**DOI:** 10.1101/2023.05.30.542868

**Authors:** Lillie M. Powell, Soo Jeon Choi, Breanna L. Haught, Ryan Demkowicz, P. Rocco LaSala, Slawomir Lukomski

## Abstract

**Background:** Increasing incidence of invasive group A *Streptococcus* (iGAS) disease has been reported in Europe and United States over the past several years. Coupled with this are observations of higher rates of resistance to non-beta lactam antimicrobials.

**Objectives:** The aim of this study was to characterize iGAS and pharyngitis isolates from West Virginia (WV), a region outside of the US national active bacteria core surveillance purview, where risk factors associated with iGAS infections are prevalent.

**Methods:** Seventy-seven invasive group A *Streptococcus* isolates were collected from sixty-seven unique patients at the J.W. Ruby Memorial Hospital Clinical Microbiology Laboratory in WV from 2021-23. Invasive isolates and twenty unique pharyngitis isolates were tested for clindamycin and erythromycin susceptibilities in the clinical laboratory. Patient demographic and clinical information was retrieved from patient electronic health records. Isolates were further characterized based on *emm*-type and detection of MLS_B_ resistance determinants.

**Results:** Twenty-six (39%) isolates were of a single *emm*-type, *emm92*. All *emm92* isolates were uniformly erythromycin/clindamycin resistant with inducible or constitutive MLS_B_ resistance imparted by the plasmid-borne *erm*(T) gene. The majority of *emm92* infections were associated with adult patients who reported intravenous drug use, whereas no pharyngitis infections were caused by an *emm92* strain. Overall, fifty-one (76%) of the sixty-seven iGAS isolates were determined to carry MLS_B_ resistance.

**Conclusions:** Isolates of *emm*-type 92 predominated in this collection, were uniformly erythromycin/clindamycin resistant, and were associated with adult intravenous drug use but not with pediatric pharyngitis.

## Introduction

Group A *Streptococcus* (GAS) is a recognized culprit of skin and soft tissue infection (SSTI), with necrotizing fasciitis, systemic dissemination, toxic shock, and high mortality representing the far end of its invasive clinical spectrum. Although GAS remains uniformly susceptible to penicillin, recommended therapeutic regimens for severe invasive group A *Streptococcus* (iGAS) infections include clindamycin in combination with a β-lactam^1^. The Centers for Disease Control and Prevention (CDC) designated erythromycin resistant GAS as a concerning threat in 2019 following observations of significant increases in both erythromycin and clindamycin resistance from U.S. surveillance sites between 2006-2017.^2^ Erythromycin resistance methylase (*erm*) genes acquired by GAS include *erm*(A), *erm*(B), and *erm*(T), all of which impart resistance to erythromycin and clindamycin.^2-5^ The *erm-*encoded enzymes methylate an A2058 residue in 23S ribosomal rRNA, thereby blocking the active binding site for all MLS_B_ (macrolide-lincosamide-streptogramin B) antibiotics.

In addition to its observation of increasing rates of resistance, the CDC antibacterial core program’s surveillance across 10 U.S. sites also identified intravenous drug use (IVDU) and homelessness as important risk factors for iGAS infections, with associations noted among particular iGAS *emm*-types.^3, 6^ The population of West Virginia (WV) has one of the highest rates of drug abuse in the U.S.^7^ but is beyond the CDC’s surveillance catchment area.^2, 3^ To counter this knowledge gap, we previously characterized 76 iGAS isolates from WV (January 2020-June 2021) and found that the majority of infections were caused by isolates harboring MLS_B_ resistance, many of which belonged to *emm*-type 92.^8^ Our findings corroborated CDC data in highlighting the emergence of *emm92-*type iGAS across the U.S. since 2010.^2, 3, 5, 8^ Therefore, considering that the WV population represents an important high-risk population outside of the current CDC surveillance area, the aim of this study was to expand the categorization of WV-iGAS isolates by examining a collection of contemporary clinical isolates producing infections between July 2021 to January 2023.

## Methods

### Isolate collection

Clinical isolates were collected from West Virginia University Medicine (WVUmed) system hospitals and retrieved from the clinical microbiology laboratory at the primary reference facility at J.W. Ruby Memorial Hospital in Morgantown, WV. The collection included all viable primary and referred iGAS isolates available from the freezer bank. A sampling of pharyngitis isolates obtained from throat swab specimens were also retrieved. Isolates spanned the period of July 2021 to January 2023 (Four isolates were collected in January 2023).

### Epidemiological data

Patient records were reviewed to capture demographic and clinical information following approval by Institutional Review Board (Protocol #2211685003), including: patient age, sex, residence location/status, history of IVDU, and infection source. All results were reported and retrieved from patient electronic health records (Epic, Verona, WI).

### Susceptibility testing

Erythromycin, clindamycin, and D-test susceptibility testing and all quality control measures were performed by disc diffusion according to the methods described by the Clinical Laboratory Standards Institute (CLSI M100-S31).^9^ Zone diameters were interpreted using CLSI clinical breakpoints, and any degree of clindamycin zone flattening in proximity to erythromycin disc was interpreted as a positive D-test result.

### Detection of resistance determinants and emm typing

Genomic DNA was isolated per the alternate lysate protocol listed on the CDC Streptococcal Epidemiology Laboratory page at https://www.cdc.gov/streplab/groupa-strep/index.html. Genomic DNA was used to determine isolate *emm*-type by Sanger sequencing of PCR products generated with primers *emm1* and *emm2*, followed by the BLAST search on the CDC database, as described.^10^ Genomic and plasmid DNA was used for PCR amplification with primers detecting resistance determinants (*erm*(A), *erm*(B), *erm*(T), and *mefA*/*E* genes) as previously indicated (Table S1).^8, 11, 12^

## Results

A total of 77 iGAS isolates recovered from 67 patients with invasive infections from July 2021-January 2023 were investigated. Median patient age was 43 years (mean 47, range 2-79; including 3 pediatric patients aged 2, 7, and 8 years) with a slight male preponderance (63%). Infected patients resided in 22 of the 55 counties in WV and surrounding counties in PA (6), OH (1), and MD (2). The 56 in-state patients represented a wide geographical distribution, with the highest proportion (46%) from three northern counties (Harrison, Marion, and Monongalia), which is likely due their higher population densities and proximity to the main WVUMed system hospital in Morgantown. The county of residence for two patients was unknown.

Analysis showed the predominate isolate type to be *emm92* (39%, n=26), with less prevalent *emm*-types including *emm11* (16%, n=11), *emm12* (7%, n=5), *emm49* (7%, n=5), *emm83* (6%, n=4), *emm58* (4%, n=3), and *emm151* (3%, n=2). The remaining isolates were of various individual *emm*-types. Isolates of historically predominate *emm*-types, ^13-15^ which emerged since the 1980s as the main causes of iGAS infections, were notably lacking in this collection, with a single isolate each of *emm1, emm3*, and *emm28* observed.

Of 57 patients with sufficient social history documented, 33% (n=22) reported a history of IVDU, over half of whom (55%, n=12) were infected with *emm92* isolates (Figure 1A, Table S2). Further, among the 26 patients harboring *emm*92 infections, 46% (n=12) had a history of IVDU compared to 31% (n=8) who did not (six patients were not assessed due to inadequate documentation). People experiencing homeless accounted for three patients in total, two of whom were infected by *emm*92 isolates.

**Figure 1:**
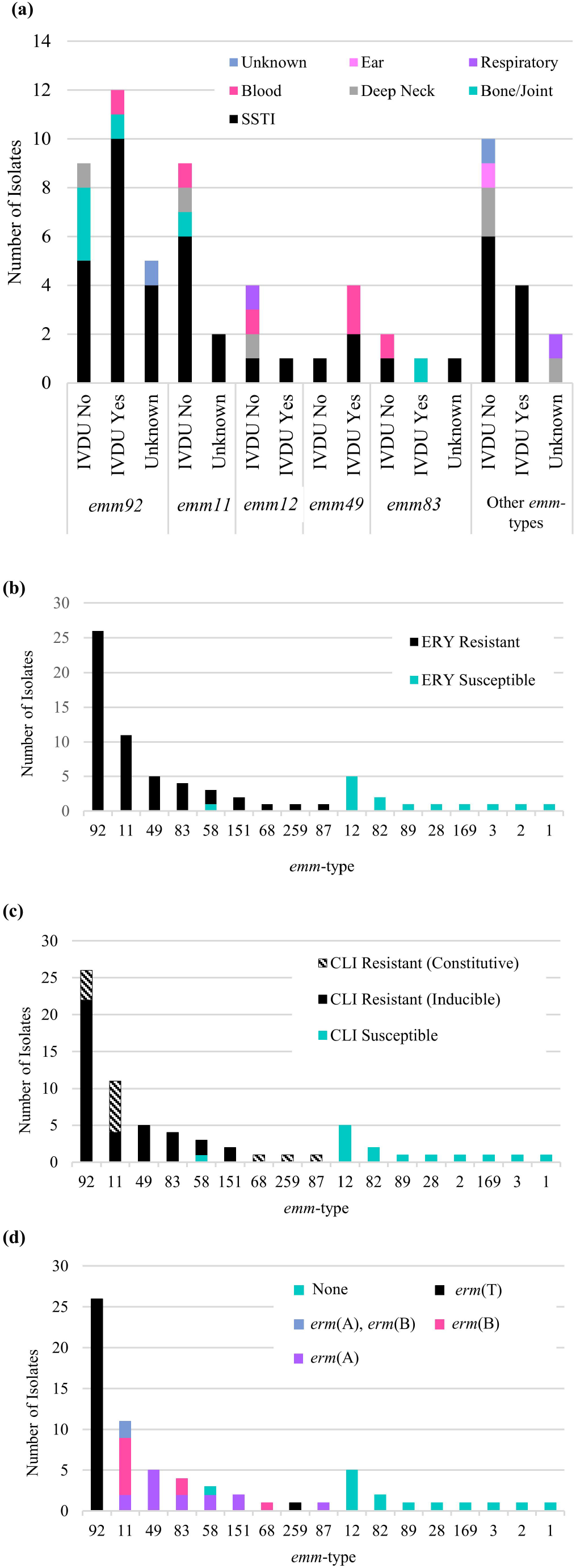
Characterization of iGAS isolates in West Virginia, USA, 2021-2023. (*a*) Assessment of *emm*-type in relation to infection source and IVDU history. *B-C*, MLS_B_ susceptibility and resistance profiles. The number of isolates resistant to erythromycin (*b*) and clindamycin (*c*) by *emm*-type. Clindamycin susceptibility or resistance was determined by D-test results where susceptible isolates did not create a zone of inhibition. Accordingly, the growth pattern in a D-test indicated if an isolate had a constitutive or inducible phenotype. *(d)* Phenotypic and genotypic characterization of resistance. Isolate genotype according to PCR-detection of resistance determinants.

Among the 67 patients, 57% (n=38) presented with an SSTI of the extremities and 7% (n=5) with SSTI of non-extremities (e.g. breast abscess, decubitis ulcer, surgical site infection). Other sources of iGAS infection included bone/joint (10%, n=7), bloodstream without a clear primary source (10%, n=7), deep neck (9%, n=6), and one each of respiratory, ear/mastoiditis, and “wound” not otherwise specified (Figure 1A, Table S2). iGAS infections resulted in the death of at least 4 patients (6%), though a substantial number of patients were discharged with limited available information on follow-up care and final outcome.

Overall, 76% (n=51) of isolates were resistant to erythromycin and clindamycin (Figure 1B, C, Table 1). In the U.S. resistance to erythromycin is encoded by either erythromycin resistance methylase genes, *erm*(A), *erm*(B), and *erm*(T), or the *mefA/E* efflux pump, the latter of which does not impart clindamycin resistance.^2-4, 16, 17^ The uniquely plasmid-borne *erm*(T) gene was detected in all 26 *emm92* isolates, the majority (85%) of which had an inducible (iMLS_B_) phenotype (i.e. clindamycin resistance detected only upon erythromycin induction). Intriguingly, both the pRW35-like plasmid harboring *erm*(T) gene and the *erm*(T)-specific PCR product were detected in a single *emm259* isolate (Figure S1) displaying constitutive (cMLS_B_) resistance (Figure 1D), which supports that other strains are capable of acquiring this mobile genetic element.^18^ By contrast, *emm11* isolates carried either *erm*(A) (n=2) or *erm*(B) (n=7) alone or in combination (n=2). Additionally, 64% of *emm11* isolates had cMLS_B_ resistance. All remaining erythromycin resistant isolates were found to carry either the *erm*(A) or *erm*(B) gene (Figure 1D), whereas no isolates with the *mefA/E* efflux pump comprising the M phenotype^16^ were detected.

**Table 1:**
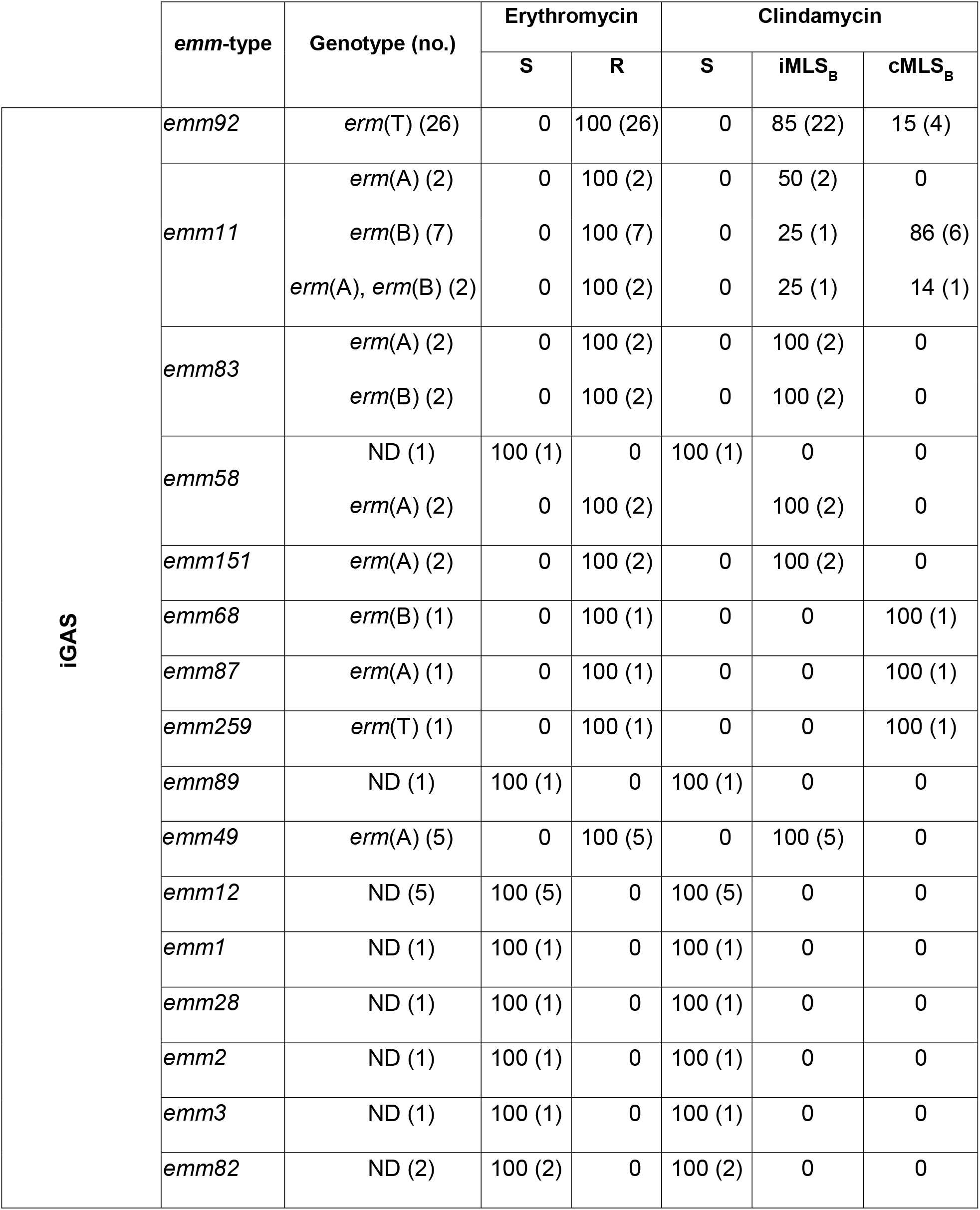
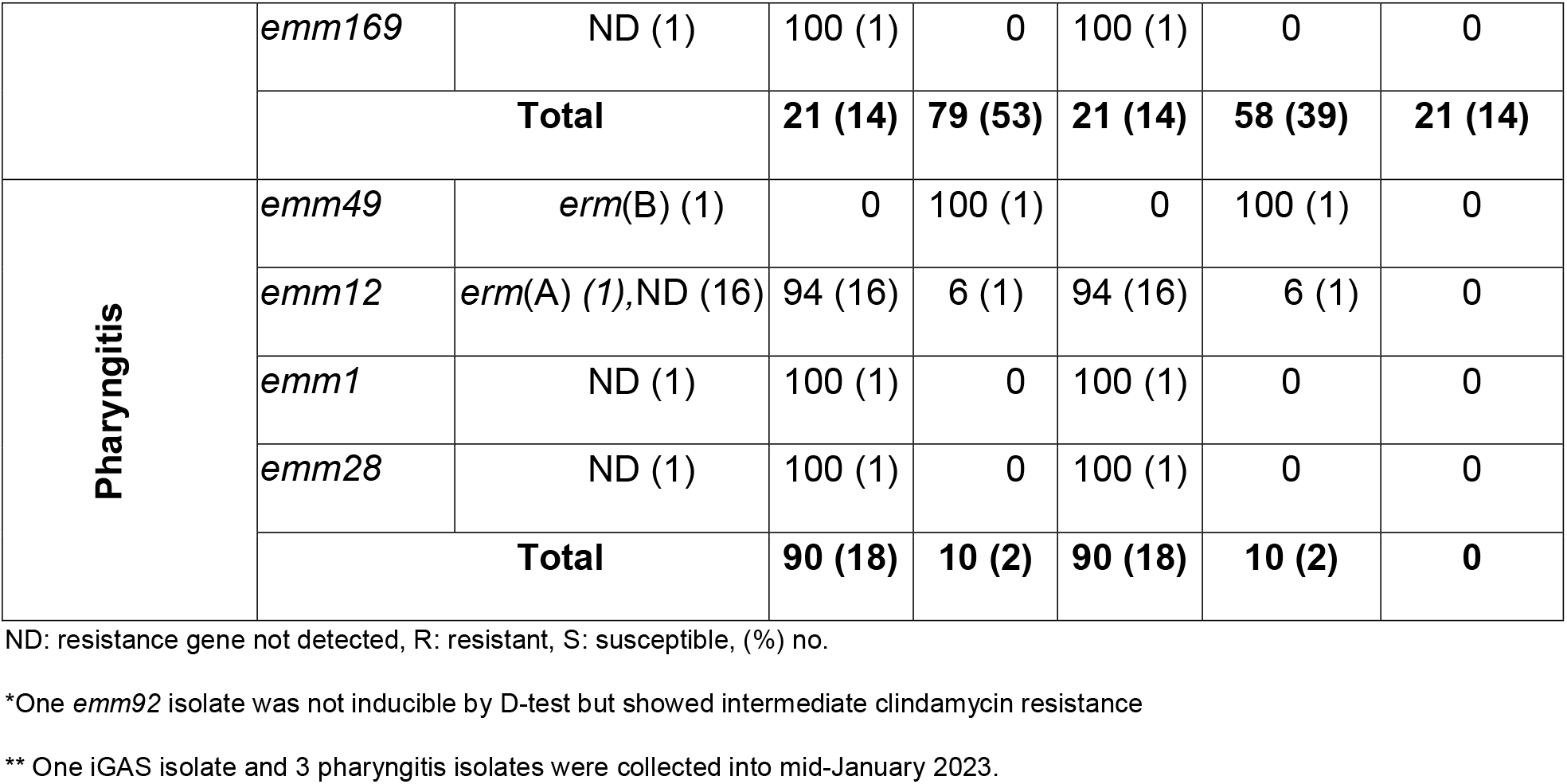
Phenotypic antimicrobial susceptibility results for iGAS and pharyngitis isolates by *emm*-type and resistance determinant listed as % (n).

The *emm92* strain is categorized as a Clade D organism, which suggests that it would be capable of causing both pharyngitis and skin infections.^13, 19^ Based on this we hypothesized that *emm92* isolates would be an important causative agent of pharyngitis. Throat isolates are not routinely banked but we collected 20 isolates for this study during the time overlapping iGAS collection. The majority (85%) of throat isolates were *emm*-type 12, as well as singular *emm1, emm28*, and *emm49* isolates. Overall, 90% of the throat isolates were susceptible to erythromycin and clindamycin. One *emm12* (erm(A)) and one *emm49* (*erm*(B)) isolate were found to have an iMLS_B_ phenotype of resistance. Altogether, this suggests that pharyngitis infections were not a major reservoir for *emm92*-type isolates in WV during the study interval.

## Discussion

This study extends previous work by our lab [8] and indicates that iGAS MLS_B_ resistance has remained alarmingly high (76% overall) in WV since 2020. Because recommendations for use of combination clindamycin and beta-lactam therapy for iGAS infections derive largely from clinical evidence that predates the widespread emergence of iMLS_B_ and cMLS_B_ phenotypes and/or that excluded patients with resistant isolates [reviewed in ^20^] we question whether such an approach to empiric treatment is warranted for all regions.

Here we also confirm the continued high prevalence of *emm92* isolates in the state’s patient population as well as a substantial association between *emm92* iGAS infections and patient history of IVDU. Taken together, the two-study collections account for 132 patient isolates recovered over a 3-year period, of which 46% were *emm92*. This proportion is considerably higher than the 15-18% noted in previous CDC data sets,^2, 3^ which may be the result of expansion of one or more sub-clonal populations within our region. Furthermore, whereas comprehensive isolate collections reveal non-MLS_B_ *emm1* as the predominant cause of iGAS infections,^2, 3, 5^ the low abundance of *emm1* isolates in WV iGAS disease (2% collectively) is a unique trend.

Importantly, *emm92*-type isolates were not associated with any pediatric pharyngitis cases investigated herein. While the small sampling of only 20 throat isolates exists as a limitation to our study, Sanson *et al*. reported the complete absence of *emm*92 among 431 pharyngitis cases in Houston, TX, despite identifying it as a cause of 11 iGAS infections.^4^ Taken together, these observations suggest pharyngitis is unlikely to be a major transmission source for adult iGAS infections by *emm*92.

In brief, this study highlights the substantial differences that may occur regionally in iGAS epidemiology and antimicrobial susceptibility rates while stimulating additional questions related to the virulence, fitness, and other attributes of emm92 that are contributing to its emergence in WV.

## Transparency declarations

All authors report no conflicts of interest.

## Acknowledgements

We would like to thank the Centers for Disease Control and Prevention’s (CDC) Emerging Infection Program laboratory for providing control GAS strains. D-test and MLS_B_ susceptibility testing were performed by the Clinical Microbiology Laboratory staff at J.W. Ruby Memorial hospital in Morgantown, WV.

## Author contributions

P.R.L., S.L., and L.P. conceived the study design and protocol. L.P. drafted the manuscript and preformed all analysis of results. P.R.L and R.D. provided clinical isolates, accessed the necessary patient data on the Epic system (according to IRB approval), and oversaw susceptibility testing performed by the Clinical Microbiology Laboratory staff at J.W. Ruby Memorial hospital in Morgantown, WV. Experimental data was collected by L.P., S.J.C., and B.H. to allow characterization for each isolate. P.R.L., R.D., and S.L. critically reviewed the manuscript. All authors reviewed the manuscript for final approval.

