## Supplemental Material for "Prevalence of Erythromycin Resistant *emm92*-type Invasive Group A Streptococcal Infections in West Virginia, United States, 2021-2023"

**Supplementary Data**

**Table S1:** Primers used for detection of *emm*-type and erythromycin (*erm*) resistance genes.

| **Gene(s)** | **Primer** | **Sequence** | **Reference** |
| --- | --- | --- | --- |
| *emm* | 1 For | 5’-TATTSGCTTAGAAAATTAA-3’ | ^1^ |
|  | 2 Rev | 5’-AAACAAGCTAAAGAACTTGC-3’ | ^1^ |
| *erm*(A)(TR) | TR For 3 | 5’-ACATCTAAAAAGCATGTAAAGG-3’ | ^2^ |
|  | TR Rev 3 | 5’-CTTCAGCACCTGTCTTAATTG-3’ | ^2^ |
| *erm*(B)(AM) | AM For | 5’-GAAAAGGTACTCAACCAAATA-3’ | ^3^ |
|  | AM Rev | 5'-AGTAACGGTACTTAAATTGTTTAC-3’ | ^3^ |
| *erm*(T) | T For | 5’-CCGCCATTGAAATAGATCCT-3’ | ^4^ |
|  | T Rev | 5’- GCTTGATAAAATTGGTTTTTGGA-3’ | ^4^ |
| *mefA/E* | A/E For | 5’-CAGTATCATTAATCACTAGTGC-3’ | ^3^ |
|  | A/E Rev | 5’-TTCTTCTGGTACTAAAAGTGG-3’ | ^3^ |

**Table S2:** Infection source according to reported patient history of IVDU and *emm*-type of patient isolates.**

| ***emm*-type** | **IVDU History** | **Source of Infection** | **Total** |
| --- | --- | --- | --- |
| ***emm92*** | **IVDU No** | SSTI | 5 |
|  |  | Bone/Joint | 3 |
|  |  | Deep Neck | 1 |
|  | **IVDU Yes** | SSTI | 10 |
|  |  | Bone/Joint | 1 |
|  |  | Blood | 1 |
|  | **Unknown** | SSTI | 4 |
|  |  | Unknown | 1 |
|  |  | **Total** | **26** |
| ***emm11*** | **IVDU No** | SSTI | 6 |
|  |  | Bone/Joint | 1 |
|  |  | Blood | 1 |
|  |  | Deep Neck | 1 |
|  | **Unknown** | SSTI | 2 |
|  |  | **Total** | **11** |
| ***emm12*** | **IVDU No** | SSTI | 1 |
|  |  | Blood | 1 |
|  |  | Deep Neck | 1 |
|  |  | Respiratory | 1 |
|  | **IVDU Yes** | SSTI | 1 |
|  |  | **Total** | **5** |
| ***emm49*** | **IVDU No** | SSTI | 1 |
|  | **IVDU Yes** | SSTI | 2 |
|  |  | Blood | 2 |
|  |  | **Total** | **5** |
| ***emm83*** | **IVDU No** | SSTI | 1 |
|  |  | Blood | 1 |
|  | **IVDU Yes** | Bone/Joint | 1 |
|  | **Unknown** | SSTI | 1 |
|  |  | **Total** | **4** |
| **All other *emm*-types** | **IVDU No** | SSTI | 6 |
|  |  | Deep Neck | 2 |
|  |  | Ear/Mastoiditis | 1 |
|  |  | Unknown | 1 |
|  | **IVDU Yes** | SSTI | 4 |
|  | **Unknown** | Deep Neck | 1 |
|  |  | Respiratory | 1 |
|  |  | **Total** | **16** |
| **Total Unique Patient Isolates** | | | **67** |
| *Ten patients had insufficient information to determine IVDU history whereas, residence status was undefined for eight patients | | | |
| **Data represented here corresponds to figure 1 panel A | | | |

**Figure S1:** Detection of plasmid DNA in resistant isolates of diverse *emm* types. The number above each lane corresponds to isolate *emm*-type. Plasmid DNA was extracted to confirm the presence of the pRW35-like plasmid, which harbors the *erm*(T) gene. Presence of the pRW35-like plasmid was confirmed in the *emm259* and *emm92* isolates.

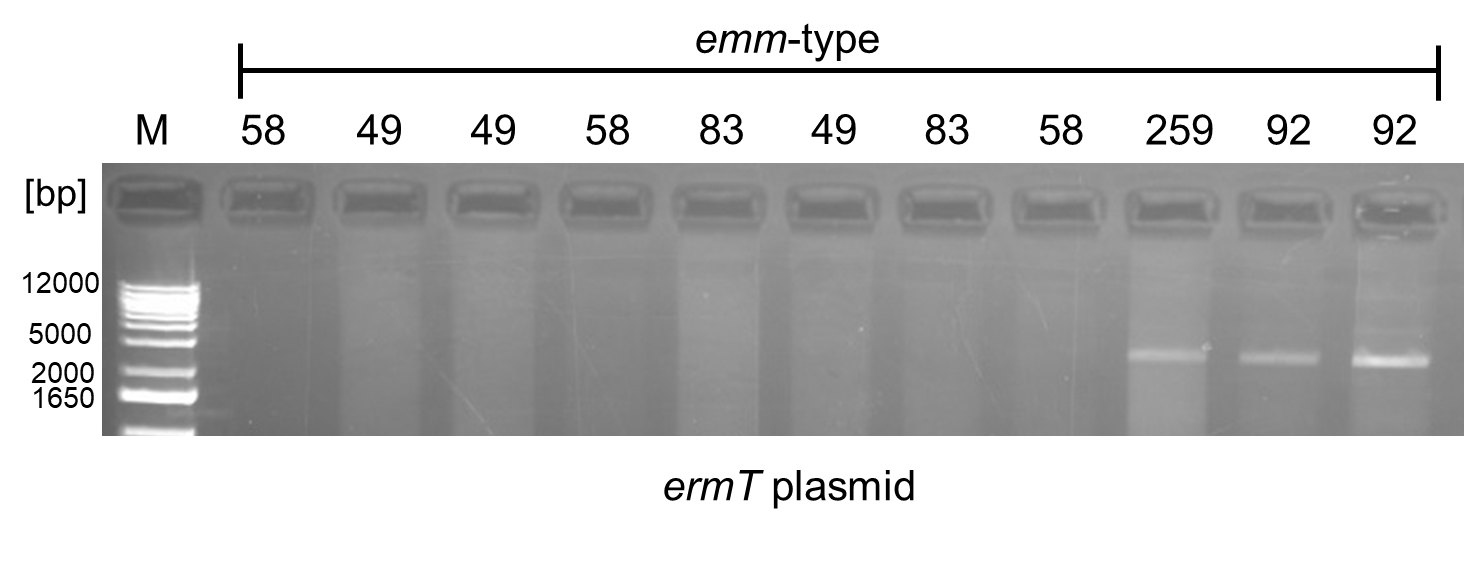
